# Genomic Analyses of Antibiotic-Resistant *Escherichia coli* From Extensive Beef Cattle and Sheep Farms Identifies Inter-Species and Farm-Farm Sharing as Clonal Dissemination Pathways

**DOI:** 10.64898/2025.12.06.692737

**Authors:** Noora Peltonen, Jordan E. Sealey, Oliver Mounsey, Caroline M. Best, Beatriz Llamazares, Will Miller, Yelyzaveta Moiseienko, Katie L. Sealey, Elliot Stanton, Emily Syvret, Lucy Vass, Laura Wright, Kristen K. Reyher, Matthew B. Avison

## Abstract

**Background:** Globally, there is a large gap in our understanding of the prevalence, ecology and transmission dynamics of antibiotic resistance (ABR) in extensively reared ruminants, despite these animals contributing to the food chain and frequently sharing land with humans.

**Methods:** Five hundred and seventy one visits to 33 Welsh beef cattle and/or sheep farms resulted in 1874 samples being collected at faecally contaminated sites from April 2022 to March 2023 (ADGC1) and September 2023 to December 2024 (ADGC2). Samples were tested for resistant *Escherichia coli* using amoxicillin, streptomycin, spectinomycin, cefotaxime and ciprofloxacin. WGS used Illumina technology. Clonal relationships were determined following core-genome alignment.

**Results:** A significant reduction in positivity for spectinomycin-resistant *E. coli* in sheep samples from ADGC1 to ADGC2 was observed, coincident with market withdrawal of a spectinomycin-containing preparation widely used in sheep. Reductions were seen in 19/22 sheep flocks with nine seeing a >50% reduction. Resistance to other tested antibiotics was unchanged. Phenotypic analysis and WGS for 713 *E. coli* showed that resistance to antibacterials important for human medicine was rare and genetically diverse. We identified 77 *E. coli* clones (<100 SNP cutoff) circulating among study farms with mixed farms contributing most; clones were also shared between animal species on mixed farms.

**Conclusions:** For extensively reared ruminants, ABR-reducing efforts can have significant impacts on antibiotic resistance on farms. Focusing these efforts onto farms contributing to the most animal movement and mixing events may generate the greatest reductions in overall on-farm ABR prevalence at regional and national levels.

## Introduction

Antimicrobials, and particularly antibiotics, are important in livestock farming, being used to support animal health, welfare and productivity. The extensive use of these medicines for decades, however, has contributed to the emergence of antibiotic-resistant (ABR) bacteria on farms.^1^ Antibiotic resistance can reduce animal welfare and food security because it increases the burden of animal disease. There is also widespread concern that ABR bacteria, and the mobile genetic elements that encode antibiotic resistance, found in livestock are entering human populations.^2–4^ This could exacerbate the global health crisis caused by ABR human infections, which has predominantly been driven by antibiotic overuse in humans.^5,6^ Faecal bacteria such as *Escherichia coli* are a likely zoonotic link between livestock and humans because (among myriad potential scenarios) faeces can contaminate the environment, including water and crops as well as meat and milk derived from livestock, and can be transferred by direct interaction between livestock and humans, their homes and workplaces.^7–13^

Small ruminants - predominantly sheep and goats farmed for wool, meat and milk - are estimated to number approximately 2 billion globally.^14^ These animals commonly range widely and are frequently traded and transported.^15,16^ Despite their global importance, very little monitoring of ABR bacteria has been undertaken in small ruminants.^17–27^ This is in contrast with more intensively farmed livestock such as pigs, poultry, feedlot beef cattle and dairy cattle.^28–30^ In the UK, 31 million sheep were recorded in 2024 compared with 9.4 million cattle and 4.7 million pigs. Yet there is a dearth of information about ABR bacteria in sheep, particularly in healthy animals sampled on farms.^24–27^

Over 90% of the land area of Wales, a devolved nation within the UK with a population of approximately 3.2 million, is used for agriculture, of which a significant proportion is used for grazing and rearing beef cattle and sheep.^31^ Most of this takes place in mountainous and hill farming areas, often unsuitable for other use.^32^ In 2024, approximately 8.75 million sheep/lambs and 1 million cattle were present on Welsh farms.^33^

In the UK, beef is typically extensive, falling into one of three categories: calf-rearer, grower-finisher or suckler. Suckler (or cow-calf) farms maintain a permanent breeding herd. Growing beef cattle may be moved between farms and systems depending on their production stage, often multiple times before slaughter.^34,35^ Despite their economic importance and regular interaction with environments frequented by people, ABR monitoring in extensive beef cattle is, like in small ruminants, very limited globally.^19,21,23–27,36–40^

As part of the Wales-wide project Arwain Defnydd Gwrthficrobaidd Cyfrifol (ADGC), we have monitored ABR *E. coli* in faecally contaminated sites around sheep and beef cattle. In this paper, we isolated and sequenced *E. coli* with a range of resistance phenotypes, including those resistant to European Medicines Agency (EMA) antibiotics categorised as Categories B (Restrict), C (Caution) and D (Prudence). This allowed us to consider whether farm-farm and inter-species sharing are associated with amplification of ABR *E. coli* clones among extensively-reared beef cattle and sheep.

## Materials and Methods

### Ethics statement

Farmers gave fully informed written consent to participate in the study. Ethical approval was obtained from both the University of Bristol’s Animal Welfare and Ethics Review Body and the Faculty of Health Sciences Research Ethics Committee (refs: VIN/21/030; 10337).

### Sample collection

A convenience sample of 33 farms was recruited in April 2022, comprising 10 sheep, 12 mixed beef/sheep and 11 beef cattle farms from locations across Wales. Basic information about the farms is presented in **Table S1**. Farms were sampled 12 times, monthly, from April 2022 until March 2023 (ADGC1) and six times, every 2-3 months, from September 2023 until December 2024 (ADGC2). There were 571 visits and 1874 samples collected: 1238 in ADGC1 and 636 in ADGC2; 976 around beef cattle and 898 around sheep. In ADGC1, we monitored positivity for *E. coli* resistant to spectinomycin in samples from sheep and streptomycin in samples from beef cattle. In ADGC2, we monitored both spectinomycin and streptomycin resistance in all samples. Resistance to amoxicillin, ciprofloxacin and cefotaxime was monitored in all samples across both projects.

A mixture of sample types was collected around animals of different ages and production stages. The objective was to give an overview of each farm. Samples were collected by a representative of the farm’s registered veterinary practice, with standard operating procedures for sampling provided. We requested that each sample contained the faeces of multiple animals. Samples were collected using sterile overshoes worn while traversing farm areas trafficked by ruminants or by spooning faeces into a sterile universal container. Samples were kept refrigerated until they were sent to the bacteriology lab using public mail services at ambient temperature. Median time from sample collection to receipt in the laboratory was 3 days with 88.7% of batches received <7 days after collection.

### Bacteriology

Approximately 1 g of each sample was weighed, and PBS added at 10 mL/g before vortexing and mixing with an equal volume of 50% sterile glycerol. Twenty microlitres of each sample was spread onto Tryptone Bile X-Glucuronide agar plates without antibiotic or containing amoxicillin (8 mg/L), spectinomycin (32 mg/L), streptomycin (32 mg/L), cefotaxime (2 mg/L) and ciprofloxacin (0.5 mg/L) to select for resistant *E. coli* as blue/green colonies. Two *E. coli* colonies per selective plate were isolated and re-streaked onto all selective agar plates to define each isolate’s antibiotic susceptibility profile. One *E. coli* isolate per antibiotic susceptibility profile per sample was considered for WGS and stored using cryobeads at -70°C. We stored isolates resistant to cefotaxime and/or ciprofloxacin from positive samples collected at Farm Visits 1 to 15 inclusive. Other resistant isolates were stored only from samples collected at Farm Visits 1, 4, 7 (ADGC1), 13, 14 and 15 (ADGC2) (**Figure S1**).

### WGS and Bioinformatics

WGS was performed on each stored isolate (**Figure S1**) by MicrobesNG using standard protocols to achieve a minimum 30-fold coverage. Genomic DNA library construction used the Nextera XT Library Prep Kit (Illumina, San Diego, USA) following the manufacturer’s protocol but with double the input DNA amount and a 45s PCR elongation time. Libraries were sequenced on an Illumina NovaSeq 6000 (Illumina, San Diego, USA) using a 250 bp paired end protocol. Reads were adapter trimmed using Trimmomatic version 0.30 ^41^ with a sliding window quality cutoff of Q15. De novo assembly used SPAdes version 3.7.^42^ Quality control was with QUAST.^43^ Exclusion criteria were: contigs >500; largest contig <250,000; total genome size <4,500,000 or >6,000,000; GC content <50.00 or >51.00. Contigs were annotated using Prokka 1.11.^44^ Fasta files were analysed using ResFinder 4.1, including pointfinder ^45^ and MLST 2.0.^46^ Sequence alignment used Snippy (4.6.0)(https://github.com/tseemann/snippy). SNP distances were determined using SNP-dists (0.8.2) (https://github.com/tseemann/snp-dists). The reference genome was PubMLST reference PDT001296075.1.

### Statistical analysis

To account for repeated sampling on farm in our analysis of ABR *E. coli* prevalence at different timepoints or types of farm, a mixed-effects logistic regression was used, including a random intercept for farm.

## Results

### Sample-level positivity for resistant E. coli

Across the 33 study farms sampled for ADGC1 (April 2022 to March 2023) and ADGC2 (September 2023 to December 2024), 11 out of 1874 collected samples were negative for *E. coli* with a limit of detection of 1000 cfu/g. Sample-level positivity for *E. coli* resistant to antibiotics representative of EMA Category B (cefotaxime, a 3^rd^-generation cephalosporin [3GC], and ciprofloxacin, a fluoroquinolone [FQ]) was <2% (**Table 1**).

**Table 1.**
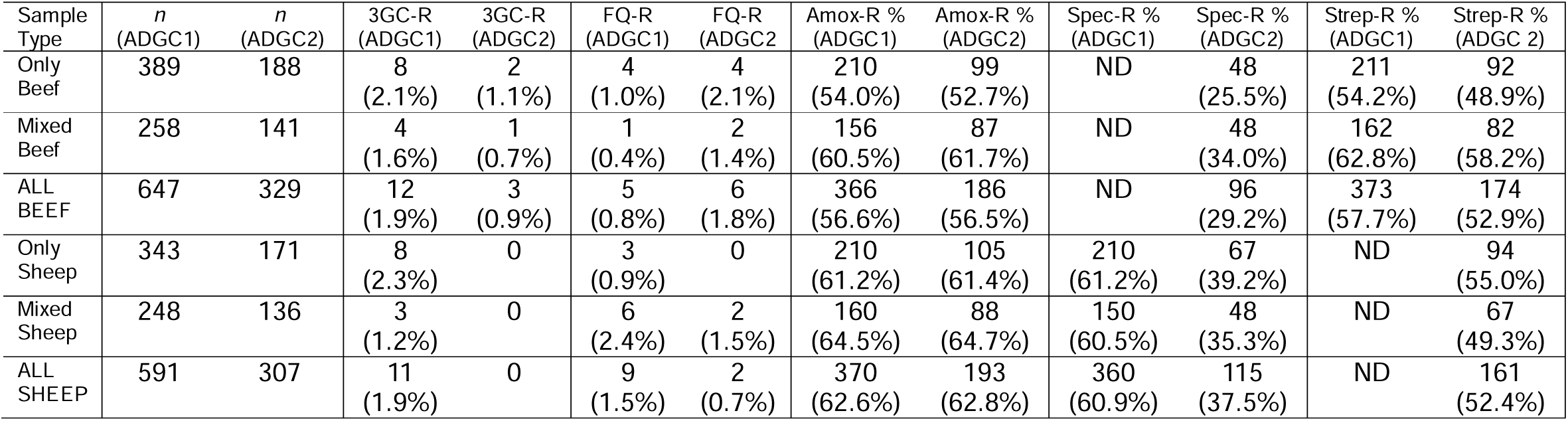
Sample-level positivity for *E. coli* resistant (-R) to test EMA Category C/D antibiotics amoxicillin (Amox), spectinomycin (Spec), streptomycin (Strep) or EMA Category B antibiotics cefotaxime (3GC) and ciprofloxacin (FQ). Data are divided by farm (only beef, mixed, only sheep), animal type (on mixed farms) and ADGC project. ND = Not Determined.

In contrast, sample-level positivity for (EMA Category C/D) amoxicillin-, streptomycin- or spectinomycin-resistant *E. coli* was high. Between ADGC1 and ADGC2 there was a significant decrease in spectinomycin-resistant *E. coli* positivity in sheep samples (Odds Ratio [OR] 0.37; 95% Confidence Interval [CI] 0.28-0.50; *p*<0.0001; **Table 1**). When considering farms individually, year-to-year fluctuation was observed for amoxicillin resistance in beef cattle and sheep as well as for streptomycin resistance in beef cattle. However, 19/22 farms showed a downward trend for spectinomycin resistance in sheep samples from ADGC1 to ADGC2, with nine farms showing a >50% reduction (**Tables S2, S3**).

In ADGC2 (where data were available for beef cattle and sheep), a trend for a greater percentage positivity for spectinomycin-resistant *E. coli* in samples was seen from sheep-only (39.2%) verses beef-only (25.5%) farms (**Table 1**; OR 2.04, 95%CI 0.86-4.85, *p*=0.11). A similar trend was not seen when comparing samples collected around beef cattle (34.0% positive) versus those collected around sheep (35.4% positive) on mixed species farms (OR 1.07, 95%CI 0.64-1.78, p=0.8).

Spectinomycin resistance is caused by the AadA aminoglycoside-modifying enzyme, which also confers resistance to streptomycin.^47^ We therefore calculated sample-level positivity for streptomycin resistance but spectinomycin susceptibility (predominantly caused by StrAB)^48^ by counting samples that were positive for streptomycin resistance but negative for spectinomycin resistance. A significantly higher percentage of beef samples fitting this profile were identified (23.7% positive) than sheep samples (15.0%; OR 0.59, 95%CI 0.38-0.91, p=0.02).

### Genomic analysis of resistance to antibiotics important for human health

**Figure S1** shows how resistant *E. coli* isolates were selected for sequencing and the numbers of isolates sequenced where data met quality control standards as well as after removing all but the first sequenced isolate from a sample with a particular sequence type (ST) and resistance gene combination. Of 713 isolates retained for analysis, 361 were from beef cattle and 352 were from sheep; 350 were from sole-species farms and 363 were from mixed-species farms. Whilst only 20 sequenced isolates came from plates containing a FQ or a 3GC (**Figure S1**), 54 unique ST/antibiotic resistance gene (ARG) combinations included ARGs relevant to antibiotics used to treat human *E. coli* infections in the UK ^49,50^ and were seen among 42/361 sequenced isolates from beef cattle and 46/352 from sheep. These combinations were spread across 23 out of the 33 study farms. Ten farms housed only one or two ST/ARG combinations, but farms M10 and M11 housed nine and seven combinations, respectively (**Tables S4, S5**).

Resistance to 3GCs was seen in 15 ST/ARG combinations with varied mechanisms including plasmid AmpCs, chromosomal AmpC hyperproduction and extended-spectrum β-lactamases, primarily CTX-M types. In only one farm (M10) was more than one 3GC resistance mechanism found. FQ resistance was seen in 11 ST/ARG combinations, generally caused by DNA gyrase/topoisomerase mutations. However, in five cases, resistance was due to the plasmid-mediated quinolone resistance mechanism QnrS1 alongside a single gyrase/topoisomerase mutation. A *qnr* gene was also found in FQ-susceptible isolates from 11 additional ST/ARG combinations. Only one isolate was both 3GC- and FQ-resistant, and this isolate also carried *bla*_OXA-484_ on an IncF plasmid (**Tables S4, S5**). This gene encodes a globally disseminated OXA-48-like carbapenemase.^51^

Five ST/ARG combinations were found on more than one study farm. An ST847 clone carrying *fosA7*, conferring resistance to fosfomycin (from EMA Category A [Avoid]), was found on 2 farms with 18 SNPs between isolates. ST1086 carrying *fosA7* was also found on two farms but the isolates differed by 629 SNPs. The most widespread fosfomycin-resistant clone was ST1086 carrying *fosA7* alongside a single mobile genetic element encoding resistance to amoxicillin, erythromycin, lincomycin, spectinomycin, streptomycin, tetracycline and sulphamethoxazole. This combination was found on six farms with pairwise SNP distances between one to 88. Overall, fosfomycin resistance was seen in 18 ST/ABR gene combinations and gentamicin resistance in six. In no case was *fosA7* found alongside resistance to any other important agents. The two other ST/ARG combinations found on multiple farms were an FQ-R ST58 clone carrying *qnrS1*, found on two farms with isolates 35 SNPs apart. Finally, FQ-R ST744 was found on two farms but isolates differed by 1461 SNPs (**Tables S4, S5**).

In addition to the ARGs reported above, 78/361 (21.6%) and 66/352 (18.8%) of sequenced *E. coli* isolates from beef cattle and sheep samples, respectively, carried trimethoprim resistance genes.

### *E. coli* clones shared between farms and between animal species

Among the 713 sequenced ABR *E. coli*, 127 and 104 STs were represented in beef and sheep samples, respectively, with 72% of isolates coming from 51 STs shared between animal species (**Table S6**). To measure in more detail pairwise relationships between sequenced isolates of these shared STs, we determined core-genome SNP distances for all possible *E. coli* isolate pairs. We identified 210 *E. coli* isolates (106 from sheep and 104 from beef cattle) clustered into 77 separate clones (**Table S7**). In this study we defined a clone as a group of isolates spanning more than one farm, where each isolate in the clone differed by <100 SNPs from at least one isolate originating on a different farm. In the absence of any agreed consensus in the field, 100 SNPs is a cutoff often used to define a clonal relationship between isolates.^52^ Where multiple isolates of a given clone were found on a farm, all except the isolate differing by the fewest SNPs from an isolate originating on a different farm were excluded (112 isolates in total), minimising the implications of repeated sampling at farm level on downstream analysis.

All 33 farms yielded at least one isolate from a clone. Thirty-one clones included isolates from more than two farms and the most populous clone was found on seven farms. There was no significant difference in the likelihood that isolates from sheep or beef cattle were found in a clone. However, there was a bias in favour of isolates collected on mixed farms being in a clone compared with those from sole-species farms (Fisher’s Exact p=0.04; **Table S7**). Plotting all study farms on a map and recording the locations of farms sharing the six most widely distributed clones (**Figure 1**) revealed dissemination over a wide geographical range.

**Figure 1.**
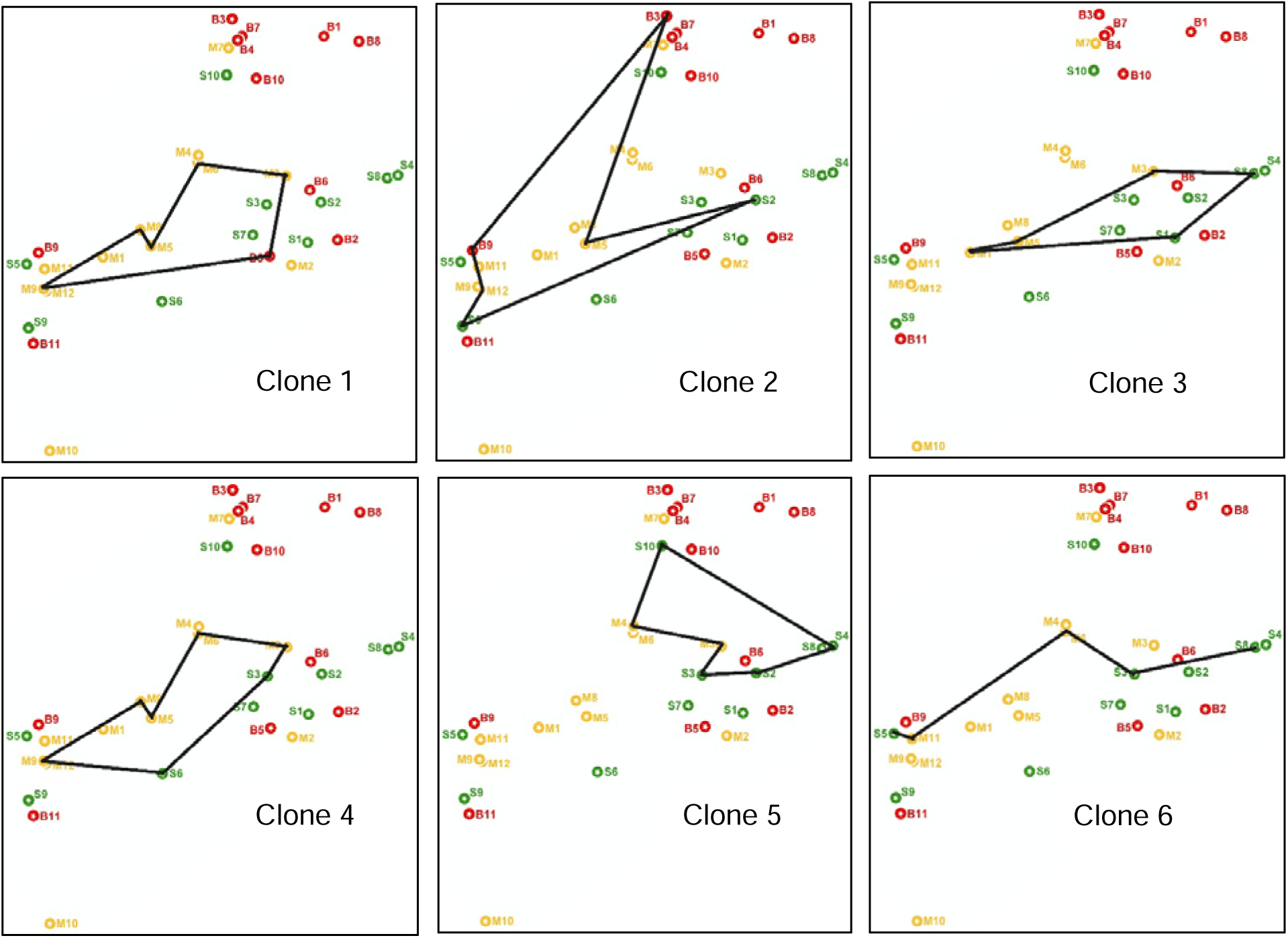
Map of study farms showing distributions of the most abundant clones. Relative positions of the study farms (red markers, B labels, beef only; green markers, S labels, sheep only; yellow markers, M labels, mixed beef/sheep farms). Farm numbers are as used throughout this report. The positions of these farms represent relative distance, but the map has undergone multiple transformations to minimise the chances of farm identification. The scale from left to right in each map is 200 km. Each map represents the “range” of each of the most abundant clones (in terms of number of farms on which the each *E. coli* clone was observed (**Table S7**). Lines merely link the farms on which these clones were found and do not imply direct transmission or directionality.

We only sequenced resistant *E. coli* isolates in this study, so the likelihood of isolates on a farm being observed to contribute to a clone may have been associated with the resistance rate on that farm. Indeed the total number of isolates sequenced on a farm was somewhat correlated with the total number of clones to which that farm contributed (R^2^=0.68). However, the ratio of clones contributed to per isolate sequenced ranged from 0.75:1 to 0.11:1 across the study farms (**Table S8**). There was some impact of farm type on this ratio. The median ratio for beef, sheep and mixed farms was 0.29, 0.38 and 0.40, respectively; Mann-Whitney, *p*=0.04 for a beef/mixed farm comparison. Sheep farms were not significantly different from the other two types.

We also identified 25 instances (involving 11/12 mixed farms) where a resistant *E. coli* clone was found in samples from sheep and beef cattle on the same farm. These pairs tended to have very low SNP distances (**Table S9**).

## Discussion

Extensively reared sheep and beef cattle are economically and culturally important in many countries, particularly because they can utilise land that is otherwise suboptimal.^32^ These extensive systems are generally considered to have low antimicrobial usage compared with more intensive farming systems; nonetheless, extensively reared ruminants have widespread interaction with human populations, primarily through shared land use.^53^ Accordingly, it is important to monitor ABR faecal bacteria such as *E. coli*, which readily contaminate the environment, in these animals. Through doing so, it will be possible to include these animals within plans designed to reduce the burden of resistant opportunistic infections in people.

This multi-year study of ABR *E. coli* on sheep and beef farms is the first large-scale attempt to address gaps in our understanding of the prevalence of ABR *E. coli* in samples collected from environments around extensively reared sheep and beef cattle. We identified high sample-level positivity for EMA Category C and D agents (50-65% in ADGC1; **Table 1**). It is not appropriate to compare positivity rates in other studies conducted by other researchers ^17–27,36–40^ because bacteriological methodology differences can have dramatic effects on observed prevalence. However, using an identical methodology,^54^ we reported positivity rates for amoxicillin-resistant *E. coli* from dairy cattle in South West England (2017/18 calendar years) to be 66%, compared with 57% in beef cattle and 63% in sheep in ADGC1 (2022-24). In contrast, for 3GC-R *E. coli*, sample-level positivity was 9.3% on English dairy farms ^54^ versus 1.9% each for beef cattle and sheep in ADGC1. For FQ-R *E. coli*, rates were 6.3.% on English dairy farms ^54^ versus 0.8% (beef) and 1.5% (sheep) in ADGC1. We should be cautious in interpreting these apparently lower rates of EMA Category B antibiotic resistance in ADGC1 beef and sheep samples compared with dairy samples, however, because we have shown that the age and location of the sample population affects sample-level positivity on dairy farms.^54^ Furthermore, since our English dairy farm study, there have been significant attempts to reduce EMA Category B antibiotic usage on farms in the UK,^55^ so we should wait for a contemporaneous survey of dairy farms and detailed epidemiological analysis before we can confirm these apparent differences between our UK dairy and beef/sheep farm surveys.

Spectinomycin is an agent that has historically been used to treat Watery Mouth Disease, an *E. coli* infection in newborn lambs that can have devastating impacts on the flock.^56^ Accordingly, we only monitored spectinomycin resistance in samples from sheep during ADGC1. We observed a significant reduction in sample-level positivity for spectinomycin-resistant *E. coli* in sheep samples between ADGC1 and ADGC2 (**Table 1**). It is notable, in the context of these findings, that spectinomycin usage in sheep across the UK is likely to have reduced from late 2021 because of the withdrawal of Spectam Scour Halt,^57^ which remains the only licenced spectinomycin-containing product for sheep in the UK.^58^ However, we have reported that, across the 22 sheep flocks studied here, spectinomycin usage was already low in 2021,^59^ perhaps reflecting veterinary prescribing behaviour changes that contributed to the decision to withdraw Spectam Scour Halt from the market.^58^ Accordingly, even if spectinomycin usage reduction has driven a decline in spectinomycin resistance rates in sheep, it may not have declined quickly, and the role of market withdrawal is unclear.

Given increased interest in spectinomycin resistance, we began measuring rates in samples from beef cattle in ADGC2. We observed a trend, though falling below the threshold required for statistical significance, towards lower spectinomycin resistance rates in beef cattle than sheep, and particularly when considering beef-only versus sheep-only farms (**Table 1**). In hindsight, it would have been beneficial to monitor spectinomycin resistance in beef cattle during ADGC1, since rates were even higher in sheep at that time (**Table 1**), so we hypothesise that the difference between cattle and sheep sample-level positivity would have been greater than what was seen in ADGC2, providing more power for our comparison. Whilst all samples have been stored frozen, we did not return to the beef samples and re-test for spectinomycin resistance because we have reported previously that freezing cattle faecal samples can result in a decline in sample-level positivity for resistant *E. coli* after around 12 months.^60^ Hence we could not be sure that this would not confound our comparison. Further sampling to increase the power of this analysis may provide more evidence that spectinomycin resistance rates are higher in samples from sheep than from cattle, potentially due to historically greater spectinomycin usage in sheep than in cattle.

Interestingly, spectinomycin resistance rates in beef cattle and sheep samples from mixed species farms were almost identical (**Table 1**). This may reflect sharing of bacteria, spreading the impact of spectinomycin usage on resistance across both species. Indeed, our WGS analysis revealed extensive evidence that ABR *E. coli* are shared between animal types on mixed farms (**Table S9**). We conclude, therefore, that mixed farms form a focus for transmission of resistant *E. coli* between animal species, potentially enhancing the circulation of these bacteria more widely.

We observed evidence of farm-farm sharing for multiple ABR *E. coli* clones (**Table S7**). Despite similar numbers of isolates being sequenced from mixed farms and sole-species farms, and after normalising for the numbers of isolates sequenced on each farm, we observed that mixed farms were more likely than sole-species farms - and particularly beef cattle farms - to contribute to ABR *E. coli* clones shared with at least one additional study farm. We take these findings concerning farm-farm and inter-species sharing as evidence for the hypothesis that stock movement and mixing (likely to be greater on mixed-species farms) are associated with *E. coli* dissemination between farms. If this hypothesis is expanded and tested through more detailed research, prime mitigation intervention points where driving down resistance could be targeted for maximum regional benefit might be identified.

## Acknowledgements

The authors would like to thank all farmers who participated in the study and veterinarians who assisted with farm recruitment and collected all samples. The authors are grateful to members of the wider Arwain DGC consortium for support and useful discussion. We particularly thank Iechyd Da (Gwledig) Limited and Mentera who assisted with veterinary practice and farmer recruitment.

## Authors’ contributions

KKR, MBA designed the study and obtained funding. CMB, LV, MBA managed the study and participant recruitment. NP, JES, BL, OM, WM, YM, KLS, ESy, LW processed samples, undertook the bacteriology and prepared isolates for whole genome sequencing. LV, CMB, ESt cleaned and curated the antibiotic resistance data. NP, JES, OM, BL, MBA performed the genomic analyses, statistical analyses, geographical mapping and data visualisation. NP, MBA wrote the manuscript. All authors read and approved the final manuscript.

