## Supplementary material for "Genomic Analyses of Antibiotic-Resistant *Escherichia coli* From Extensive Beef Cattle and Sheep Farms Identifies Inter-Species and Farm-Farm Sharing as Clonal Dissemination Pathways": Tables S1-S9 and Figure S1

**[2]Bristol Veterinary School, Langford House, Langford, Bristol, BS40 5DU, United Kingdom**

**[†]Corresponding Author**

**Table S1. Demographic data for the study farms**

| | Sheep flocks (n = 22) | | Beef herds (n = 23) | |
|---|---|---|---|---|
| | **Sheep-only farms (n = 10)** | **Mixed beef & sheep farms (n = 12)** | **Beef-only farms (n = 11)** | **Mixed beef & sheep farms (n= 12)** |
| **Land size (hectares)** | | | | |
| Mean | 112.7 | 240.0 | 113.0 | 240.0 |
| Minimum | 22 | 59 | 41 | 59 |
| Maximum | 445 | 526 | 304 | 688 |
| **Herd size[1]** | | | | |
| Mean | - | - | 161.3 | 153.8 |
| Minimum | - | - | 63 | 30 |
| Maximum | - | - | 274 | 400 |
| **Flock size[2]** | | | | |
| Mean | 535.2 | 776.3 | - | - |
| Minimum | 99 | 205 | - | - |
| Maximum | 1100 | 2750 | - | - |
| **Lambing location** | | | | |
| Majority indoors | 6 (60%) | 6 (50%) | - | - |
| Majority outdoors | 4 (40%) | 6 (50%) | - | - |
| **Stratification system[3]** | | | | |
| Lowland | 6 (60%) | 3 (25%) | - | - |
| Upland | 5 (50%) | 7 (58%) | - | - |
| Hill | 1 (10%) | 5 (42%) | - | - |
| **Organic status** | | | | |
| Non-organic (conventional) | 10 (100%) | 12 (100%) | 10 (91%) | 11 (100%) |
| Organic | 0 (0%) | 0 (0%) | 1 (9%) | 0 (0%) |
| **Enterprise type[4]** | | | | |
| Calf rearing unit | - | - | 2 (18%) | 3 (25%) |
| Suckler herd | - | - | 8 (73%) | 8 (67%) |
| Growing unit | - | - | 5 (45%) | 4 (33%) |
| Finishing unit | - | - | 3 (27%) | 4 (33%) |

[1] Total number of cattle (all ages) recorded on farm

[2] Total number of adult breeding ewes recorded on farm

[3] Sheep farms could be comprised of multiple stratification system tiers

[4] Individual beef farms could be comprised of multiple enterprise types

**Table S2. Sample-level positivity for EMA Category C/D antibiotics in samples collected around beef cattle on beef only (B) or mixed beef & sheep farms (M).**

| Farm Code | NUMBER OF SAMPLES COLLECTED | | SAMPLES WITH AMOXICILLIN RESISTANCE (%) | | SAMPLES WITH STREPTOMYCIN RESISTANCE (%) | | |
|---|---|---|---|---|---|---|---|
| | ADGC 1 | ADGC 2 | ADGC 1 | ADGC 2 | ADGC 1 | ADGC 2 | KEY |
| B1 | 16 | 18 | 69% | 39% | 67% | 33% | -10-29% |
| B2 | 36 | 18 | 21% | 22% | 33% | 28% | -30-49% |
| B3 | 33 | 18 | 75% | 56% | 69% | 50% | -50-69% |
| B4 | 36 | 18 | 53% | 39% | 53% | 28% | ->70% |
| B5 | 36 | 18 | 72% | 83% | 56% | 72% | +10-29% |
| B6 | 36 | 18 | 78% | 89% | 78% | 94% | +30-49% |
| B7 | 36 | 18 | 67% | 50% | 61% | 28% | +50-69% |
| B8 | 22 | 18 | 44% | 72% | 61% | 44% | +>70% |
| B9 | 36 | 19 | 50% | 68% | 66% | 68% | |
| B10 | 22 | 16 | 50% | 13% | 28% | 56% | |
| B11 | 24 | 9 | 9% | 33% | 18% | 22% | |
| M1 | 16 | 12 | 69% | 83% | 69% | 58% | |
| M2 | 24 | 10 | 41% | 90% | 55% | 90% | |
| M3 | 36 | 12 | 68% | 42% | 73% | 33% | |
| M4 | 38 | 12 | 54% | 58% | 58% | 83% | |
| M5 | 24 | 12 | 81% | 75% | 75% | 67% | |
| M6 | 32 | 12 | 79% | 83% | 63% | 67% | |
| M7 | 16 | 12 | 38% | 58% | 67% | 50% | |
| M8 | 34 | 12 | 44% | 58% | 38% | 67% | |
| M9 | 26 | 12 | 85% | 67% | 92% | 67% | |
| M10 | 21 | 10 | 48% | 80% | 48% | 60% | |
| M11 | 24 | 14 | 75% | 36% | 63% | 50% | |
| M12 | 23 | 11 | 43% | 18% | 48% | 9% | |

**Coloured shading represents the percentage rise (orange) or fall (green) of sample-level positivity between ADGC1 and 2.**

**Table S3. Sample-level positivity for EMA Category C/D antibiotics in samples collected around sheep on sheep only (S) or mixed beef & sheep (M) farms.**

| Farm Code | NUMBER OF SAMPLES COLLECTED | | SAMPLES WITH AMOXICILLIN RESISTANCE (%) | | SAMPLES WITH SPECTINOMYCIN RESISTANCE (%) | | |
|---|---|---|---|---|---|---|---|
| | ADGC 1 | ADGC 2 | ADGC 1 | ADGC 2 | ADGC 1 | ADGC 2 | KEY |
| S1 | 30 | 18 | 63% | 56% | 67% | 50% | -10-29% |
| S2 | 36 | 18 | 64% | 83% | 78% | 78% | -30-49% |
| S3 | 36 | 18 | 75% | 83% | 50% | 39% | -50-69% |
| S4 | 36 | 18 | 67% | 61% | 67% | 22% | ->70% |
| S5 | 36 | 17 | 69% | 47% | 67% | 53% | +10-29% |
| S6 | 32 | 18 | 47% | 72% | 53% | 33% | +30-49% |
| S7 | 36 | 18 | 64% | 44% | 58% | 17% | +50-69% |
| S8 | 29 | 17 | 41% | 47% | 38% | 18% | +>70% |
| S9 | 37 | 12 | 62% | 50% | 59% | 17% | |
| S10 | 35 | 17 | 54% | 65% | 71% | 59% | |
| M1 | 16 | 12 | 75% | 42% | 31% | 17% | |
| M2 | 22 | 14 | 36% | 71% | 45% | 7% | |
| M3 | 26 | 12 | 88% | 75% | 73% | 75% | |
| M4 | 24 | 12 | 58% | 100% | 58% | 75% | |
| M5 | 16 | 12 | 63% | 25% | 56% | 17% | |
| M6 | 24 | 12 | 75% | 100% | 75% | 42% | |
| M7 | 24 | 12 | 50% | 75% | 38% | 25% | |
| M8 | 16 | 12 | 63% | 42% | 69% | 50% | |
| M9 | 20 | 10 | 75% | 50% | 80% | 40% | |
| M10 | 24 | 9 | 67% | 89% | 71% | 44% | |
| M11 | 17 | 8 | 65% | 50% | 71% | 13% | |
| M12 | 19 | 11 | 58% | 55% | 53% | 18% | |

**Coloured shading represents the percentage rise (orange) or fall (green) of sample-level positivity between ADGC1 and 2.**

| Isolate(s) | Farm | ST | Fluoroquinolones (B) | 3GCs (B) | Gentamicin (C) | Fosfomycin (A) | EMA Category C/D |
|---|---|---|---|---|---|---|---|
| 237052 | B2 | 69 | | | *aac(3)-IV* | | *bla*$_{TEM-1}$, *strAB, tetA, floR* |
| 236973 (2) | B3 | 154 | | *bla*$_{CTX-M-55}$ | | | |
| 247064 (2) | B3 | 58 | | *bla*$_{CTX-M-55}$ | | | *bla*$_{TEM-1}$, *strAB, aph(3')-Ia, tetB, sul2* |
| 236974 | B3 | 69 | | *bla*$_{CTX-M-55}$ | | | *bla*$_{TEM-1}$, *strAB, tetA, sul1, dfrA7* |
| 236972 | B3 | 1086[††]** | | | | *fosA7* | *bla*$_{CARB-2}$, *aadA2b, ereB, tetB, sul1* |
| 236962 | B6 | 10668 | *qnrB4* | *bla*$_{DHA-1}$ | | | *bla*$_{TEM-1}$, *strAB, tetA, mphA, sul1, sul2, dfrA17* |
| 277054 | B7 | 68 | | *bla*$_{CMY-2}$ | | | |
| 265892 | B8 | 398 | *qnrS1* | | | | *bla*$_{TEM-1}$, *strAB, tetA, floR, sul2* |
| 232424 | B8 | 3889 | *qnrS1* | | | | *bla*$_{TEM-1}$, *aadA1, strAB, aph(3')-Ia, tetA, tetM, cmlA1, floR, sul3, dfrA12* |
| 236979 | B8 | 1421 | *gyrA* S83L, *parC* S80I | | | | *bla*$_{TEM-135}$, *tetA* |
| 276978 | B9 | 10 | *qnrS1* | | | | *bla*$_{TEM-135}$, *bla*$_{CARB-2}$, *aadA2b, aph(3')-Ia, tetB, floR, ereB, sul1, dfrA36* |
| 245129 (2) | B9 | 1086[††]** | | | | *fosA7* | *bla*$_{CARB-2}$, *aadA2b, ereB, tetB, sul1* |
| 237065 | B10 | 642 | | *ampC* -42C>T | | | |
| 236978 | M1 | 744[††]** | *gyrA* S83L, D87N *parC* A56T, S80I | | | | *bla*$_{TEM-1}$, *aadA5, strAB, tetB, catA1, mphA, sul1, sul2, dfrA17* |
| 237155 | M1 | 1086[††]** | | | | *fosA7* | *strAB, tetB, sul2* |
| 265899 | M4 | 56 | | *ampC* -42C>T | | | *strAB, tetB, sul2* |
| 276325 | M5 | 2723 | | | | *fosA7* | *strAB, sul2* |
| 265926 | M5 | 2723 | | | | *fosA7* | *strAB, msrE, sul2* |
| 237059 (4) | M5 | 1086[††]** | | | | *fosA7* | *bla*$_{CARB-2}$, *aadA2b, ereB, tetB, sul1* |
| 265927 | M5 | 2723 | | | | *fosA7* | *strAB, tetX6, sul2* |
| 276326 | M6 | 1140 | | | *ant(2'')-Ia* | | *aadA1, strAB, aph(3')-Ia, floR, tetA, sul2, dfrA36* |
| 265940 | M8 | 1140 | | | *ant(2'')-Ia* | | *aadA1, strAB, floR, tetA, sul2, dfrA36* |
| 245162 | M8 | 847[††]** | | | | *fosA7* | *bla*$_{TEM-1}$, *strAB, tetB* |
| 265906 (2) | M10 | 58[††]** | *qnrS1* | | | | *bla*$_{TEM-1}$, *tetA, dfrA14* |
| 276974 | M10 | 10 | *gyrA* S83L, D87N *parC* S80I | | | | *bla*$_{TEM-1}$, *aadA2, strAB, tetA, floR, mphA, sul2, sul3, dfrA12* |
| 276987 (2) | M10 | 206[††] | *qnrS1, parC* A56T | | | | *bla*$_{OXA-10}$, *aadA1, tetA, cmlA1, floR, dfrA14* |
| 245169 | M10 | 3234[††] | | | | *fosA7* | *bla*$_{TEM-1}$, *strAB, tetB* |
| 268015 | M11 | 14241 | | | | *fosA7* | *aadA1, tetA, sul1* |
| 237069 | M11 | 1086[††]** | | | | *fosA7* | *bla*$_{CARB-2}$, *aadA2b, ereB, tetB, sul1* |
| 277081 | M11 | 1086 | | | | *fosA7* | *bla*$_{CARB-2}$, *aadA2b, tetB, sul1* |
| 268006 | M11 | 1086 | | | | *fosA7* | *bla*$_{OXA-1}$, *bla*$_{TEM-30}$, *bla*$_{CARB-2}$, *aadA2b, strAB, ereB, tetB, sul1, dfrA36* |
| 265932 | M11 | 342 | | | | *fosA7* | *strAB, sul2* |
| 245164 (2) | M12 | 1086[††]** | | | | *fosA7* | *bla*$_{CARB-2}$, *aadA2b, ereB, tetB, sul1* |

**Table S4. Genomic analysis of *E. coli* resistant to antibiotics used to treat *E. coli* infections in humans (EMA category in brackets) which were found in faecal samples collected around beef cattle. **ST/ABR gene combination found on multiple farms; [††]ST/ABR combination also found in samples collected around beef cattle.**

| Isolate(s) | Farm | ST | Fluoroquinolones (B) | 3GCs (B) | Gentamicin (C) | Fosfomycin (A) | EMA Category C/D |
|---|---|---|---|---|---|---|---|
| 283281 | S1 | 58[††] | *qnrS1* | | | | $bla_{TEM-1}$, *tetA*, *dfrA14* |
| 265870 | S1 | 362 | | | *aac(3)-IId* | | $bla_{TEM-1}$, *aadA1*, *strAB*, *mphA*, *tetA*, *sul1*, *sul2* |
| 237127 | S2 | 17 | *qnrB4* | $bla_{DHA-1}$ | | | *mphA*, *sul1*, *dfrA17* |
| 283291 (2) | S2 | 58 | *qnrS1* | | | | $bla_{TEM-1}$, *aadA22*, *tetA*, *dfrA14* |
| 265962 | S2 | 10 | *qnrB4* | $bla_{DHA-1}$ | | | $bla_{TEM-1}$, *aadA1*, *strAB*, *mphA*, *msrE*, *sul1*, *sul2*, *dfrA1* |
| 237140 (2) | S2 | 1086[††]** | | | | *fosA7* | $bla_{CARB-2}$, *aadA2b*, *ereB*, *tetB*, *sul1* |
| 237153 (3) | S4 | 58 | | *ampC* -42C>T | | | |
| 237131 (6) | S5 | 58 | *qnrS1* | $bla_{CTX-M-15}$ | | | |
| 237149 | S7 | 744[††]** | *gyrA* S83L, D87N *parC* A56T, S80I | | | | $bla_{TEM-1}$, *aadA5*, *strAB*, *mphA*, *catA1*, *tetB*, *sul1*, *sul2*, *dfrA17* |
| 246946 | S7 | 206 | *qnrS1*, *parC* A56T | | | | *aadA1*, *aadA2*, *cmlA*, *tetA*, *sul3*, *dfrA12* |
| 232131 (2) | S7 | 38 | | | *ant(2'')-Ia* | | *aadA1*, *floR*, *sul1*, *sul2*, *dfrA36* |
| 237147 | S9 | 744 | *gyrA* S83L, D87N *parC* A56T, S80I | | | | *aadA5*, *strAB*, *catA1*, *tetB*, *sul1*, *sul2*, *dfrA17* |
| 246974 | S9 | 14707 | | | | *fosA7* | $bla_{CARB-2}$, *aadA2b*, *tetB*, *sul1* |
| 283356 | S9 | 847 | | | | *fosA7* | *aadA1*, *tetA*, *sul1* |
| 232138 | M2 | 1086[††]** | | | | *fosA7* | *strAB*, *tetB*, *sul2* |
| 276408 (2) | M4 | 155 | *qnrS1* | | | | $bla_{TEM-176}$, *aph(3')-Ia*, *tetA*, *floR*, *dfrA14* |
| 247079 | M4 | 744 | *gyrA* S83L, D87N *parC* A56T, S80I | | | | $bla_{TEM-1}$, *aadA5*, *strAB*, *tetB*, *sul1*, *sul2*, *dfrA17* |
| 246926 | M4 | 10 | *qnrS1* | | | | $bla_{TEM-1}$, *aadA2*, *aph(3')-Ia*, *strAB*, *tetA*, *sul1*, *sul2*, *dfrA12* |
| 246977 | M4 | 746 | | $blaT_{EM-52}$ | | | *tetA* |
| 246904 | M5 | 847[††]** | | | | *fosA7* | $bla_{TEM-1}$, *strAB*, *tetB* |
| 232138 (2) | M5 | 1086[††]** | | | | *fosA7* | $bla_{CARB-2}$, *aadA2b*, *ereB*, *tetB*, *sul1* |
| 246905 | M8 | 847[††]** | | | | *fosA7* | $bla_{TEM-1}$, *strAB*, *tetB* |
| 283344 | M9 | 342 | | | | *fosA7* | *strAB*, *tetB*, *sul2* |
| 247078 | M10 | 155 | *qnrS1*, *gyrA* S83A | | | | $bla_{TEM-1}$, *strAB*, *tetA*, *sul2* |
| 237144 | M10 | 744 | *gyrA* S83L, D87N *parC* A56T, S80I | | | | $bla_{TEM-1}$, *aph(3')-Ia*, *strAB*, *catA1*, *tetB*, *sul2* |
| 237137 | M10 | 540 | *qnrS1*, *gyrA* S83L | $bla_{CTX-M-15}$ | | | $bla_{OXA-484}$, *mphA*, *tetB* |
| 283347 (2) | M10 | 206[††] | *qnrS1*, *parC* A56T | | | | $bla_{OXA-10}$, *aadA1*, *cmlA*, *floR*, *tetA*, *dfrA14* |
| 246979 | M10 | 1795 | | *ampC* -42C>T | | | |
| 232399 | M10 | 3234[††]** | | | | *fosA7* | $bla_{TEM-1}$, *strAB*, *tetB* |
| 237145 | M11 | 744 | *gyrA* S83L, D87N *parC* A56T, S80I | | | | $bla_{TEM-1}$, *aadA5*, *aph(3')-Ia*, *strAB*, *mphA*, *catA1*, *tetB*, *sul1*, *sul2*, *dfrA17* |
| 237134 | M11 | 155 | | *ampC* -42C>T | | | *aadA1*, *tetB* |
| 265960 | M11 | 1086[††]** | | | | *fosA7* | $bla_{CARB-2}$, *aadA2b*, *ereB*, *tetB*, *sul1* |
| 246936 | M12 | 33 | | | *aac(3)-IId* | | *aadA22* |

**Table S5. Genomic analysis of *E. coli* resistant to antibiotics used to treat *E. coli* infections in humans (EMA category in brackets) which were found in faecal samples collected around sheep. **ST/ABR gene combination found on multiple farms; [††]ST/ABR combination also found in samples collected around beef cattle.**

| ST | BEEF SAMPLES | SHEEP SAMPLES |
|---|---|---|
| 58 | 31 | 31 |
| 10 | 22 | 27 |
| 155 | 22 | 40 |
| 201 | 18 | 8 |
| 69 | 15 | 13 |
| 1086 | 15 | 13 |
| 56 | 12 | 2 |
| 362 | 9 | 13 |
| 101 | 8 | 3 |
| 154 | 7 | 3 |
| 641 | 7 | 2 |
| 2522 | 7 | 1 |
| 446 | 5 | 1 |
| 14696 | 3 | 9 |
| 162 | 4 | 8 |
| 1084 | 4 | 8 |
| 57 | 1 | 7 |
| 43 | 2 | 6 |
| 297 | 3 | 5 |
| 394 | 1 | 5 |
| 744 | 1 | 5 |
| 206 | 2 | 4 |
| 661 | 4 | 3 |
| 8103 | 3 | 3 |
| 949 | 2 | 3 |
| 457 | 1 | 3 |
| 847 | 1 | 3 |
| 973 | 1 | 3 |
| 1629 | 1 | 3 |
| 278 | 3 | 2 |
| 111 | 2 | 2 |
| 224 | 2 | 2 |
| 1131 | 2 | 2 |
| 1125 | 1 | 2 |
| 1722 | 1 | 2 |
| 88 | 4 | 1 |
| 5082 | 4 | 1 |
| 337 | 2 | 1 |
| 753 | 2 | 1 |
| 117 | 1 | 1 |
| 118 | 1 | 1 |
| 196 | 1 | 1 |
| 342 | 1 | 1 |
| 348 | 1 | 1 |
| 349 | 1 | 1 |
| 2175 | 1 | 1 |
| 2853 | 1 | 1 |
| 3234 | 1 | 1 |
| 4198 | 1 | 1 |
| 7096 | 1 | 1 |
| 8185 | 1 | 1 |
| BEEF SPECIFIC | 115 | |
| SHEEP SPECIFIC | | 82 |
| TOTAL | 361 | 352 |

**Table S6. ST breakdown of sequenced *E. coli* isolates from samples collected around beef cattle and sheep.**

| Isolate | Farm/ Type | Clonal Isolate(s) /Farm/ TYPE (SNP) | | | | | |
|---|---|---|---|---|---|---|---|
| 232163 | B4 BEEF | 276320/B10/BEEF(17) | 237182/B1/BEEF (22) | | | | |
| 232165 | B7 BEEF | 277000/B5/BEEF (28) | 232406/B11/BEEF (36) | 247071/M4/MBEEF (36) | | | |
| 232166 | B3 BEEF | 232426/M7/MBEEF (15) | 245158/M3/MBEEF (39) | 232434/B8/BEEF (47) | | | |
| 232167 | B5 BEEF | 283343/M9/MSHEEP (9) | 268013/M8/MBEEF (12) | 246928/M5/MSHEEP (14) | 276400/M3/MSHEEP (15) | 237203/M6/MSHEEP (25) | |
| 232414 | B3 BEEF | 245120/B4/BEEF (18) | | | | | |
| 232416 | B5 BEEF | 232432/M11/MBEEF (33) | | | | | |
| 232425 | B9 BEEF | 246903/M4/MSHEEP (36) | | | | | |
| 232438 | B3 BEEF | 232396/S3/SHEEP (14) | 246898/S1/SHEEP (18) | | | | |
| 232440 | B6 BEEF | 246964/S3/SHEEP (21) | 277053/B3/BEEF (71) | | | | |
| 236961 | B6 BEEF | 245175/M6/MBEEF (70) | | | | | |
| 236972 | B3 BEEF | 245165/M12/MBEEF (0) | 237069/M11/MBEEF (1) | 237059/M5/MBEEF (25),<br>237201/M5/MSHEEP (25) | 245129/B9/BEEF (25) | 246974/S9/SHEEP (28) | 237140/S2/SHEEP (29) |
| 236974 | B3 BEEF | 283274/S9/SHEEP (75) | 237114/M3/MSHEEP (83) | | | | |
| 237052 | B2 BEEF | 272209/M4/MBEEF (29) | 232150/M12/MBEEF (37) | | | | |
| 245121 | B5 BEEF | 265870/S1/SHEEP (11) | | | | | |
| 245126 | B7 BEEF | 246906/M8/MSHEEP (18) | | | | | |
| 245127 | B9 BEEF | 276425/M7/MSHEEP (29) | | | | | |
| 249530 | B9 BEEF | 237133/M11/MSHEEP (5)<br>236971/M11/MBEEF (7) | | | | | |
| 265901 | B9 BEEF | 265921/B10/BEEF (0) | 283342/M9/MSHEEP (79) | | | | |
| 268035 | B8 BEEF | 276994/B1/BEEF (44) | | | | | |
| 265922 | B10 BEEF | 283285/S1/SHEEP (11) | 237197/M3/MSHEEP (22) | | | | |
| 268044 | B9 BEEF | 245172/M9/MBEEF (46) | | | | | |
| 276310 | B5 BEEF | 276338/B10/BEEF (0) | | | | | |
| 276346 | B5 BEEF | 237185/S1/SHEEP (38) | 237173/M5/MSHEEP (55) | | | | |
| 276995 | B2 BEEF | 268040/M5/MBEEF (12) | | | | | |
| 277004 | B6 BEEF | 246899/S3/SHEEP (5) | | | | | |
| 277011 | B9 BEEF | 283346/M9/MSHEEP (83) | | | | | |
| 277036 | B4 BEEF | 277052/B2/BEEF (28) | | | | | |
| 283178 | B4 BEEF | 277073/M6/MBEEF (17)<br>276412/M6/MSHEEP (19) | 246972/S8/SHEEP (21) | | | | |
| 232141 | M6 MBEEF | 276394/S3/SHEEP (18) | | | | | |
| 232148 | M3 MBEEF | 283305/M4/MSHEEP (77) | 283279/M1/MBEEF (98) | | | | |
| 232160 | M8 MBEEF | 283294/S3/SHEEP (17) | 268020/M12/MBEEF (22) | | | | |
| 232404 | M6 MBEEF | 246927/M4/MSHEEP (9) | | | | | |
| 232411 | M11 MBEEF | 246900/S5/SHEEP (93) | | | | | |
| 232423 | M6 MBEEF | 236966/M8/MBEEF (0) | 232431/M10/MBEEF (15) | | | | |
| 232429 | M9 MBEEF | 236969/M10/MBEEF (0) | | | | | |

| | | | | | | | |
|---|---|---|---|---|---|---|---|
| 232430 | M10 MBEEF | 236970/M11/MBEEF (1) | | | | | |
| 236957 | M1 MBEEF | 232395/S1/SHEEP (29) | 246913/S8/SHEEP (30) | 277019/M5/MBEEF (33) | 283301/M3/MSHEEP (34) | | |
| 236963 | M2 MBEEF | 283345/M9/MSHEEP (37) | | | | | |
| 236967 | M9 MBEEF | 283289/S2/SHEEP (79) | 232146/M7/MSHEEP (85) | 276989/M11/MBEEF (87) | | | |
| 236978 | M1 MBEEF | 237147/S9/SHEEP (6) | | | | | |
| 237061 | M8 MBEEF | 237121/S3/SHEEP (58) | 246958/M9/MSHEEP (66) | 283312/M6/MSHEEP (69) | 258396/M5/MBEEF (73) | 283321/S6/SHEEP (74) | 283300/M3/MSHEEP (83) |
| 237112 | M3 MBEEF | 279657/M4/MBEEF (3) | 237122/S3/SHEEP (14) | 265930/M8/MBEEF (54) | | | |
| 245162 | M8 MBEEF | 246904/M5/MSHEEP (18) | | | | | |
| 245167 | M11 MBEEF | 246907/M9/MSHEEP (7) | 246917/S10/SHEEP (8) | | | | |
| 245178 | M4 MBEEF | 268025/M8/MBEEF (26) | | | | | |
| 265896 | M1 MBEEF | 237130/M5/MSHEEP (16) | | | | | |
| 265906 | M10 MBEEF | 265961/S2/SHEEP (24) | 283281/S1/SHEEP (35) | | | | |
| 265929 | M8 MBEEF | 283296/S3/SHEEP (23) | | | | | |
| 265931 | M10 MBEEF | 283298/M2/MSHEEP (28) | 246914/S8/SHEEP (35) | | | | |
| 268023 | M3 MBEEF | 237172/M4/MSHEEP (0)<br>268024/M4/MBEEF (0) | 237179/S2/SHEEP (3) | 246949/S10/SHEEP (14) | 237198/S4/SHEEP (16) | 237191/S3/SHEEP (42) | |
| 268117 | M6 MBEEF | 237220/S9/SHEEP (14) | | | | | |
| 272207 | M3 MBEEF | 237117/M9/MSHEEP (40) | 272223/M10/MSHEEP (61) | | | | |
| 272211 | M6 MBEEF | 246937/S1/SHEEP (6) | | | | | |
| 272212 | M7 MBEEF | 246968/S6/SHEEP (65) | | | | | |
| 276300 | M1 MBEEF | 246901/S7/SHEEP (34) | 277058/M2/MBEEF (48) | | | | |
| 276302 | M7 MBEEF | 246975/S10/SHEEP (16) | | | | | |
| 276324 | M4 MBEEF | 276339/M1/MBEEF (0) | | | | | |
| 276329 | M10 MBEEF | 283334/M7/MSHEEP (23) | | | | | |
| 276343 | M9 MBEEF | 276344/M10/MBEEF (1) | | | | | |
| 276983 | M5 MBEEF | 272217/M3/MSHEEP (31) | | | | | |
| 277085 | M7 MBEEF | 283339/M8/MSHEEP (26) | | | | | |
| 232140 | M6 MSHEEP | 276409/M4/MSHEEP (3) | | | | | |
| 232149 | M12 MSHEEP | 232132/S3/SHEEP (36) | | | | | |
| 237115 | M8 MSHEEP | 246894/M2/MSHEEP (4) | | | | | |
| 237194 | M2 MSHEEP | 276392/S2/SHEEP (59) | 237211/S7/SHEEP (82) | | | | |
| 237202 | M6 MSHEEP | 237204/S5/SHEEP (6) | 237218/M11/MSHEEP (6) | 237192/S3/SHEEP (10) | 246947/S8/SHEEP (10) | | |
| 237213 | M8 MSHEEP | 237206/S6/SHEEP (23) | | | | | |
| 246902 | M1 MSHEEP | 276430/S8/SHEEP (54) | | | | | |
| 276399 | M3 MSHEEP | 246912/S7/SHEEP (94) | | | | | |
| 276408 | M4 MSHEEP | 237142/S9/SHEEP (57) | | | | | |
| 276410 | M4 MSHEEP | 237223/S10/SHEEP (47) | | | | | |
| 276414 | M6 MSHEEP | 276446/S10/SHEEP (0) | | | | | |
| 237188 | S2 MSHEEP | 237224/S10/SHEEP (27) | 237207/S6/SHEEP (34) | | | | |

| | | | |
|---|---|---|---|
| 276416 | M6 MSHEEP | 283333/M7/MSHEEP (5) | |
| 276436 | M10 MSHEEP | 237124/S4/SHEEP (27) | |
| 232136 | S5 SHEEP | 246915/S9/SHEEP (14) | |
| 232397 | S7 SHEEP | 246967/S5/SHEEP (16) | 246918/S10/SHEEP (19) |

**Table S7. *E. coli* clones spanning two or more study farms sharing <100 SNPs between isolate pairs. The type of farm/sample type is noted: MSHEEP and MBEEF refer to samples from sheep or beef cattle from mixed farms. Numbers in brackets report the SNP distance from the index isolate (column 1). Shading indicates close relationships <20 SNPs between isolates.**

| Farm | No of Shared clones | No of sequenced Isolates | Clones per sequenced isolate ratio |
|---|---|---|---|
| B3 | 6 | 8 | 0.75 |
| S1 | 7 | 10 | 0.70 |
| S3 | 11 | 17 | 0.65 |
| S8 | 5 | 9 | 0.56 |
| M3 | 12 | 23 | 0.52 |
| M6 | 14 | 27 | 0.52 |
| S10 | 7 | 15 | 0.47 |
| M7 | 8 | 18 | 0.44 |
| M8 | 12 | 27 | 0.44 |
| M5 | 10 | 23 | 0.43 |
| M4 | 13 | 30 | 0.43 |
| B10 | 4 | 10 | 0.40 |
| B5 | 6 | 16 | 0.38 |
| S6 | 4 | 11 | 0.36 |
| S9 | 6 | 17 | 0.35 |
| B4 | 4 | 12 | 0.33 |
| S2 | 6 | 18 | 0.33 |
| M1 | 7 | 22 | 0.32 |
| M9 | 11 | 35 | 0.31 |
| M12 | 4 | 13 | 0.31 |
| B2 | 3 | 10 | 0.30 |
| M11 | 9 | 30 | 0.30 |
| B9 | 7 | 24 | 0.29 |
| B11 | 1 | 4 | 0.25 |
| S5 | 4 | 16 | 0.25 |
| M10 | 9 | 40 | 0.23 |
| B8 | 2 | 9 | 0.22 |
| S7 | 4 | 19 | 0.21 |
| M2 | 5 | 25 | 0.20 |
| B1 | 2 | 12 | 0.17 |
| B7 | 2 | 12 | 0.17 |
| B6 | 3 | 23 | 0.13 |
| S4 | 2 | 19 | 0.11 |

**Table S8. Farms ranked by the number of clones (<100 SNPs) which they participate within divided by the number of isolates sequenced, with only the first isolate per clone per farm considered in the denominator.**

| Isolate | Farm/ Type | Related Isolate(s) /Farm/ TYPE (SNP) |
|---|---|---|
| 277044 | M1 MBEEF | 283279/M1/MSHEEP (3) |
| 236957 | M1 MBEEF | 246892/M1/MSHEEP (5) |
| 265896 | M1 MBEEF | 265897/M1/MSHEEP (1) |
| 268022 | M1 MBEEF | 232157/M1/MSHEEP (1) |
| 232418 | M2MBEEF | 265951/M2/MSHEEP (2) |
| 237112 | M3 MBEEF | 237113/M3/MSHEEP (4) |
| 268024 | M4 MBEEF | 237172/M4/MSHEEP (0) |
| 237059 | M5 MBEEF | 237201M5/MSHEEP (0) |
| 276983 | M5 MBEEF | 283311/M5/MSHEEP (0) |
| 277073 | M6 MBEEF | 276412/M6/MSHEEP (2) |
| 232404 | M6MBEEF | 272225/M6/MSHEEP (3) |
| 265928 | M6 MBEEF | 265956/M6/MSHEEP (4) |
| 268117 | M6 MBEEF | 272218/M6/MSHEEP (0) |
| 277020 | M6 MBEEF | 265972/M6/MSHEEP (1) |
| 272212 | M7 MBEEF | 237174/M7/MSHEEP (7) |
| 276302 | M7 MBEEF | 276426/M7/MSHEEP (9) |
| 232160 | M8 MBEEF | 237175/M8/MSHEEP (1) |
| 237061 | M8 MBEEF | 237116/M8/MSHEEP (1) |
| 245159 | M8 MBEEF | 246931/M8/MSHEEP (1) |
| 245162 | M8 MBEEF | 246905/M8/MSHEEP (0) |
| 245169 | M10 MBEEF | 232399/M10/MSHEEP (10) |
| 276987 | M10 MBEEF | 283349/M10/MSHEEP (0) |
| 236971 | M11 MBEEF | 237133/M11/MSHEEP (2) |
| 277031 | M11 MBEEF | 283352/M11/MSHEEP (0) |
| 245161 | M12 MBEEF | 246922/M12/MSHEEP (2) |

**Table S9. Pairs of *E. coli* isolates from mixed beef/sheep farms sharing <10 SNPs where one member of the pair was from beef cattle and the other from sheep samples on the same farm. Farm codes and the sample type are noted.**

**Figure S1. Resistant *E. coli* isolates selected for sequencing and attrition following deduplication and QC analysis.**

**EMA Category B Antibiotics**

From 1564 samples (Farm Visits 1-15 inclusive), 42 ciprofloxacin and cefotaxime plates were positive.

Following antibiotic resistance phenotypic deduplication at sample level, 34 isolates were sequenced

11 genomes failed QC checks; 3 genomes were deduplicated at sample level based on same ST and ABR gene complement

**20 unique genomes remained**:

2 from Beef-Only Farms

9 from Sheep-Only Farms

2 from Beef on Mixed Farms

7 from Sheep on Mixed Farms

**EMA Category C/D Antibiotics**

From 656 samples (Farm Visits 1, 4, 7, 13,14, 15), 862 amoxicillin, spectinomycin and streptomycin plates were positive.

Following antibiotic resistance phenotypic deduplication at sample level, 911 isolates were sequenced

139 genomes failed QC checks; 79 genomes were deduplicated at sample level based on same ST and ABR gene complement

**693 unique genomes remained**:

167 from Beef-Only Farms

172 from Sheep-Only Farms

190 from Beef on Mixed Farms

164 from Sheep on Mixed Farms
